## Supplementary information for "Coordination chemogenetics for activation of GPCR-type glutamate receptors in brain tissue"

**Supplementary table 1 | Distance (Å) between α-carbon atoms estimated by the crystal structure of mGlu1 open form (PDB 1EWT) and closed form (PDB 1EWK).**

| | H54-N264 | H55-N264 | E60-N264 | H111-N264 |
|---|---|---|---|---|
| Apo | 20.1 | 16.3 | 16.8 | 17.4 |
| Glutamate binding | 13.9 | 10.2 | 10.4 | 10.9 |

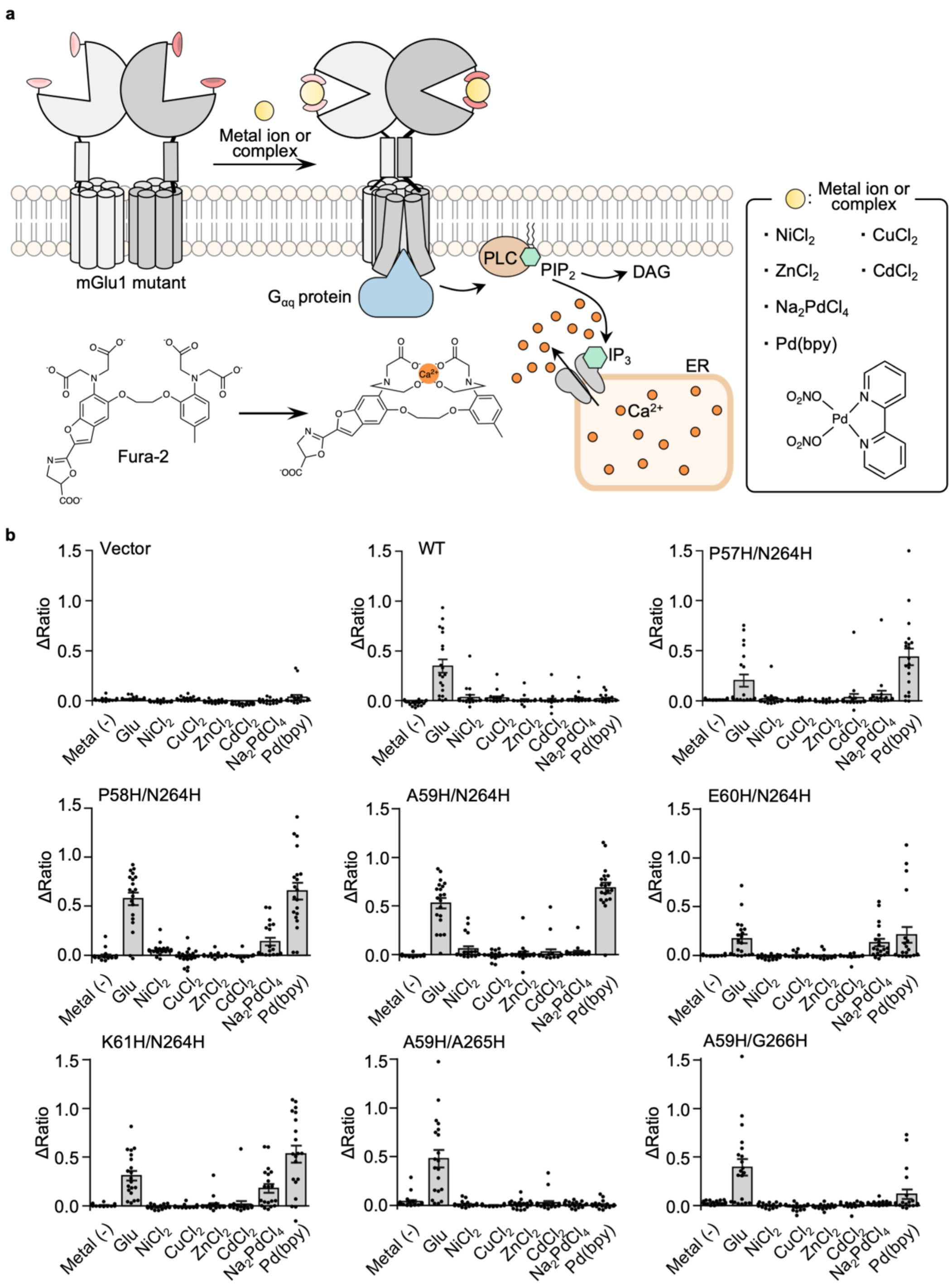


**Supplementary figure 1 | Screening assay for mGlu1 double mutants with metal ions and a metal complex. (a)** Schematic illustrations of detection of $[Ca^{2+}]_i$ changes after activation of Gq-coupled mGlu1 using Fura-2. **(b)** Averaged Δratio induced by 10 μM of glutamate, metal ions, or metal complex. Data are presented as mean ± s.e.m.

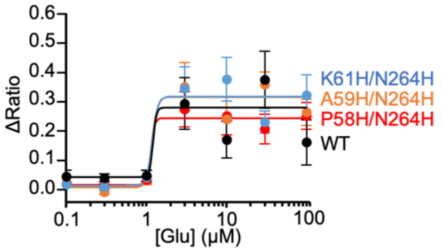


**Supplementary figure 2 | Concentration-dependent curves of glutamate for hit mutants or WT mGlu1.** Concentration-dependent curves for glutamate in HEK293 cells expressing mGlu1 WT (black), P58H/N264H (red), A59H/N264H (orange) or K61H/N264H (blue). (n = 20). Data are presented as mean ± s.e.m.

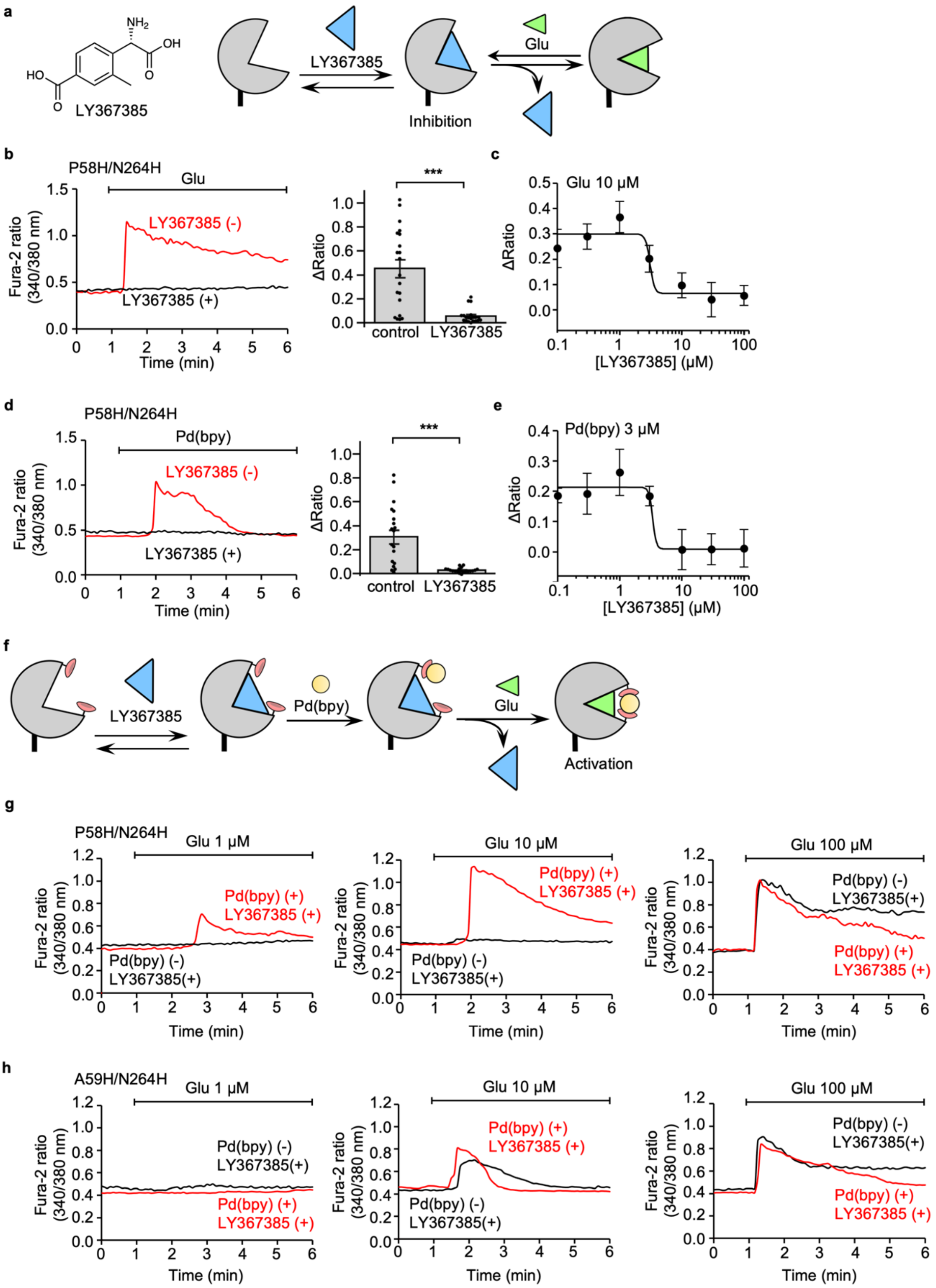


**Supplementary figure 3 | Evaluation of positive allosteric effects of the Pd(bpy) for mGlu1 mutants by co-treatment of competitive antagonist, LY367385. (a)** Schematic illustrations of competitive inhibition of mGlu1 using LY367385. (**b, c**) Inhibition of the 10 μM glutamate-induced responses using LY367385 in HEK293 cells transfected with the plasmid of mGlu1 P58H/N264H mutant. In **b**, representative traces of $Ca^{2+}$ response with (black) or without (red) 10 μM LY367385 are shown (n = 20), and averaged Δratio is shown in the right. ***Significant difference ($P < 0.001$, One-way ANOVA with Dunnet's test). In **c**, a concentration-dependent curve of LY367385 for the inhibition of 10 μM glutamate-induced response is shown. (n = 20). (**d, e**) Inhibition of the 3 μM Pd(bpy)-induced responses using LY367385 in HEK293 cells transfected with the plasmid of mGlu1 P58H/N264H mutant. In **d**, representative traces of $Ca^{2+}$ response with (black) or without (red) 10 μM LY367385 are shown (n = 20), and averaged Δratio is shown in the right. ***Significant difference ($P < 0.001$, One-way ANOVA with Dunnet's test). In **e**, a concentration-dependent curve of LY367385 for the inhibition of 3 μM Pd(bpy)-induced response is shown. (n = 20). (**f**) Schematic illustrations of the evaluation of positive allosteric effect of Pd(bpy) for glutamate-induced responses in the presence of LY367385. (**g, h**) Representative trace of glutamate-induced $Ca^{2+}$ response with (black) or without (red) 10 μM Pd(bpy) in the presence of 10 μM of LY367385 in HEK293 cells transfected with the plasmid of mGlu1 P58H/N264H (in **g**) or A59H/N264H (in **h**) mutant. Data are presented as mean ± s.e.m.

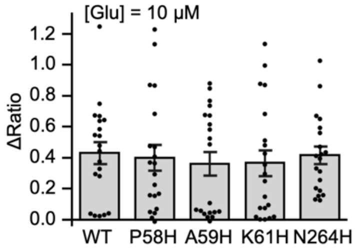


**Supplementary figure 4 | Averaged Δratio induced by 10 μM of glutamate for mGlu1 WT and single mutants.** Glutamate-induced mGlu1 responses were evaluated in HEK293 cells expressed with WT mGlu1, mGlu1(P58H), mGlu1(A59H), mGlu1(K61H), or mGlu1(N264H) mutant. (n = 20). Data are presented as mean ± s.e.m.

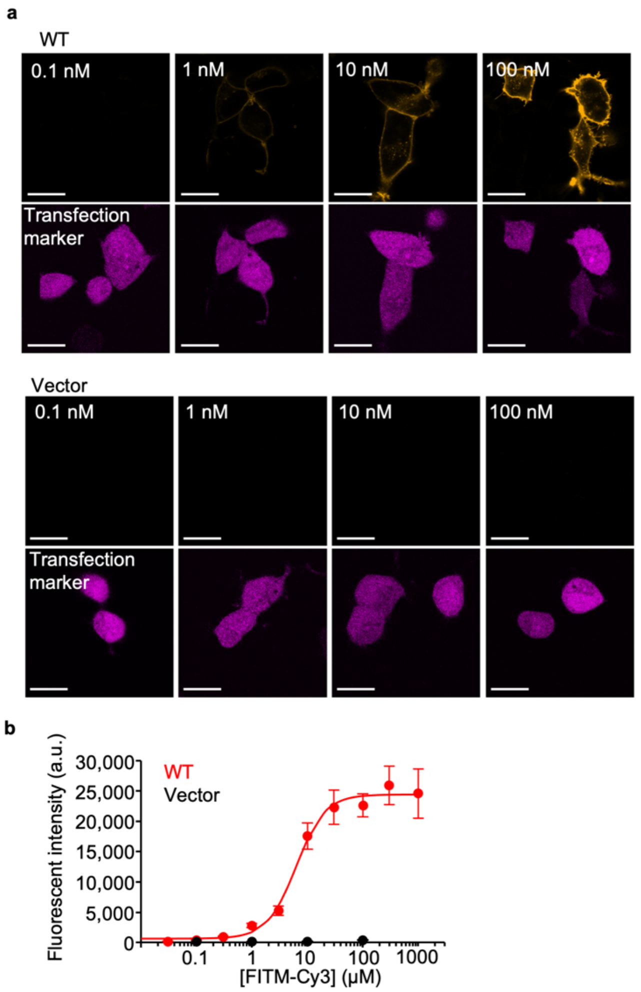


**Supplementary figure 5 | Evaluation of the affinity of FITM-Cy3 to mGlu1 WT by confocal microscopy.** (**a**) Confocal live imaging of surface mGlu1 using various concentrations of FITM-Cy3 (upper) and transfection marker (lower) in HEK293 cells expressing WT mGlu1 or the control vector. iRFP670 was utilized as a transfection marker. (**b**) Concentration-dependent curves for fluorescent intensity of FITM-Cy3 in HEK293 cells expressing WT mGlu1 (red) and vector control (black) (n= 26–118). The $K_d$ value of FITM-Cy3 to WT mGlu1 on the cell-surface was determined to be 6.8 ± 2.7 nM (n = 3, biologically independent experiments). Data are presented as mean ± s.e.m.

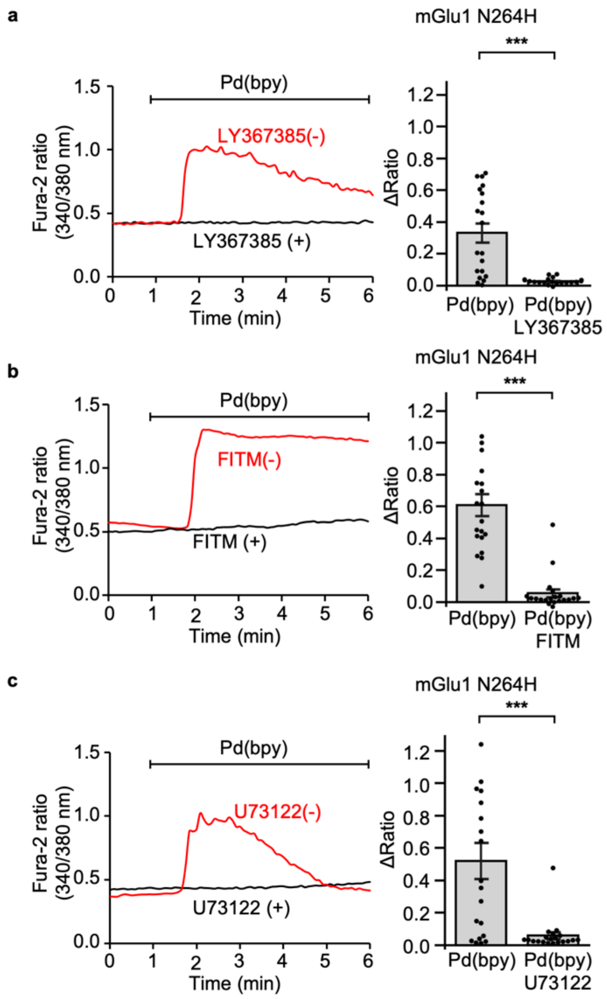


**Supplementary figure 6 | Inhibition of Pd(bpy) induced-$Ca^{2+}$ responses using LY367385, FITM, or U73122.** Left: Representative trace of 10 µM Pd(bpy)-induced responses pre-treated with 10 µM LY367385 (in **a**), 1 µM FITM (in **b**), or 10 µM U73122 (in **c**) in HEK293 cells expressing mGlu1 N264H. Right: averaged Δratio. (n = 20). ***Significant difference ($P < 0.001$, two-tailed Welch's t-test). Data are presented as mean ± s.e.m.

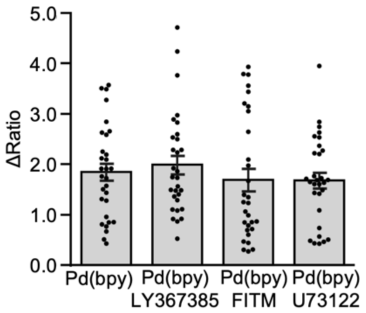


**Supplementary figure 7 | The abnormal $Ca^{2+}$ response of Pd(bpy) was inhibited by neither mGlu1 inhibitors nor a PLC inhibitor in the cultured cortical neurons.** Averaged Δratio of 30 μM Pd(bpy)-induced responses in cultured cortical neuron co-treated with 30 μM LY367385, 1 μM FITM, or 10 μM U73122. (n = 30). Data are presented as mean ± s.e.m.

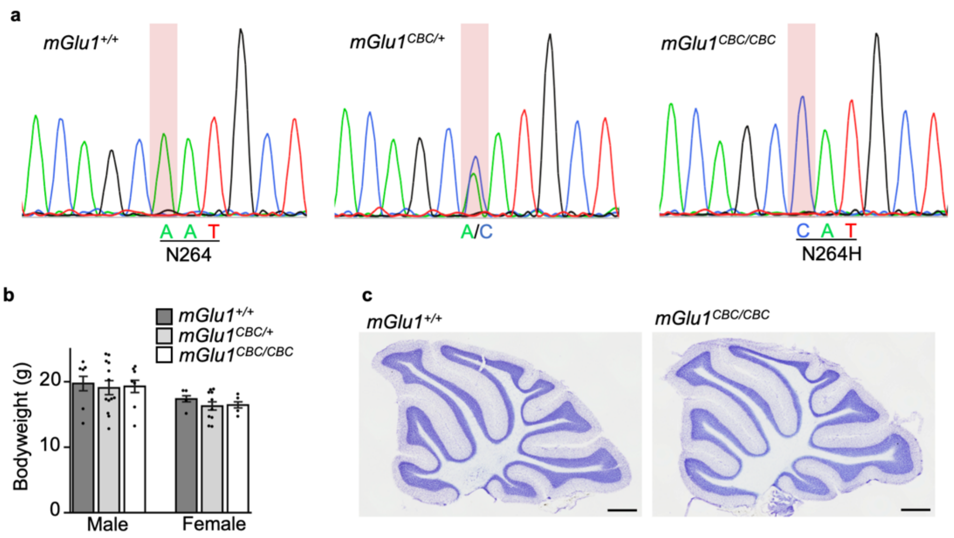


**Supplementary figure 8 | Evaluation of functionality of the mGlu1 knock-in mouse.** (**a**) Sequence analyses of exon 3 of *mGlu1* using *mGlu1*$^{+/+}$, *mGlu1*$^{CBC/+}$, or *mGlu1*$^{CBC/CBC}$ genome. (**b**) Bodyweight of the *mGlu1*$^{+/+}$, *mGlu1*$^{CBC/+}$ and *mGlu1*$^{CBC/CBC}$ mice (6 weeks old). (n= 4–13) (**c**) Cresyl violet-stained sagittal sections of adult *mGlu1*$^{+/+}$ and *mGlu1*$^{CBC/CBC}$ mouse cerebella. Scale bar, 500 µm. Data are presented as mean ± s.e.m.

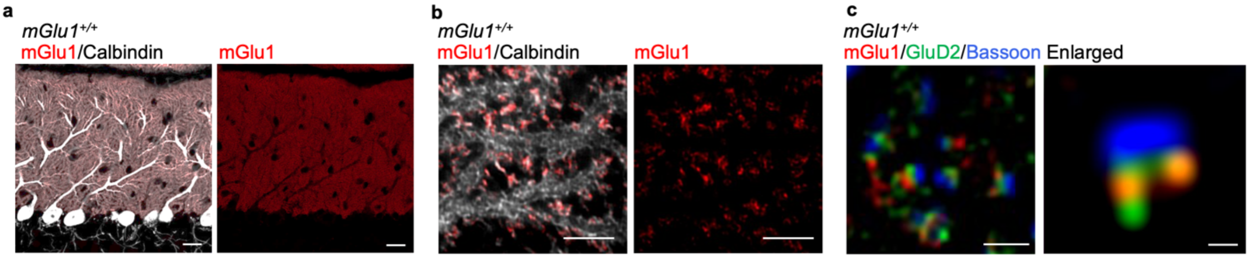


**Supplementary figure 9 | Immunohistochemistry using anti-mGlu1 antibody in *mGlu1*$^{+/+}$ cerebellar slice.** (**a**, **b**) Conventional confocal (**a**) and super-resolution (**b**) microscopy images of immune-positive signals for mGlu1 (red) and calbindin (white) in the molecular layer of a *mGlu1*$^{+/+}$ cerebellar slice. Scale bars, 20 µm (in **a**) and 2 µm (in **b**). (**c**) Super-resolution microscopic images of immune-positive signals for mGlu1 (red), GluD2 (green) and Bassoon (blue). Scale bars, 1 µm. An enlarged view is shown in a right panel. Scale bars, 200 nm.

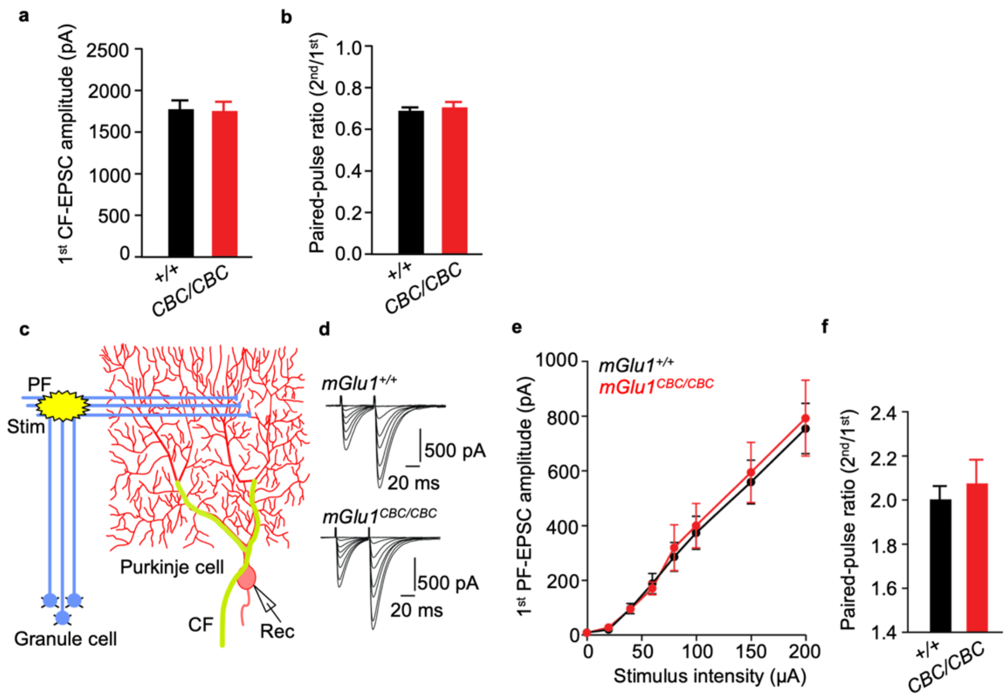


**Supplementary figure 10 | Basic synaptic transmission is normal at CF and PF–Purkinje cells in *mGlu1*$^{CBC/CBC}$ mice.** (**a**, **b**) Histograms showing CF-EPSC amplitude (**a**) and a paired-pulse ratio of CF-EPSC amplitudes (The amplitude of 2$^{nd}$ EPSC is normalized by that of 1$^{st}$ EPSC) in *mGlu1*$^{+/+}$ (+/+; black columns, n = 32 cells) and *mGlu1*$^{CBC/CBC}$ (CBC/CBC; red columns, n = 30 cells, $P > 0.05$ by Mann-Whitney U test) mice. (**c, d**) An orientation of stimulus and recording electrodes and stimulus condition (in **c**) to evoke PF-EPSCs (in **d**). In these experiments, EPSCs were evoked by the paired-pulse stimulation with various stimulus intensities (0, 20, 40, 60, 80, 100, 150, 200 μA; 50-ms inter-stimulus interval). (**e, f**) Input–Output relationship (in **e**) and paired-pulse ratio (in **f**) of the PF-EPSCs from *mGlu1*$^{+/+}$ (black, n = 13 cells) or *mGlu1*$^{CBC/CBC}$ (red, n = 13 cells, $P > 0.05$ by two-way repeated ANOVA in (**e**) and Mann-Whitney U test in (**f**)) mice. Data are represented as mean ± s.e.m.

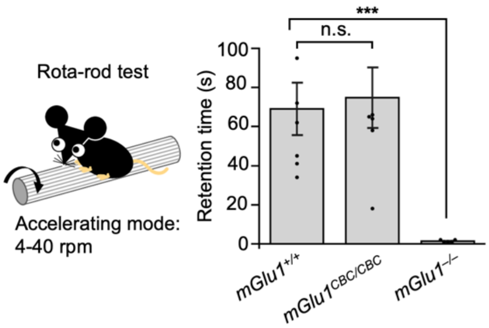


**Supplementary figure 11 | Motor coordination is normal in *mGlu1$^{CBC/CBC}$* mice.** To evaluate motor coordination of the mouse, accelerating rotor-rod test (4–40 rpm for 5 min) was performed. Histogram shows averaged retention time on rota-rod in each mouse groups. High performance was observed in *mGlu1$^{+/+}$* and *mGlu1$^{CBC/CBC}$* mice but not *mGlu1$^{-/-}$* mice. (n = 5–7). ***, $P < 0.001$ and ns, $P > 0.05$ by Kruskal-Wallis test followed by the Scheffe *post hoc* test. Data are presented as mean ± s.e.m.

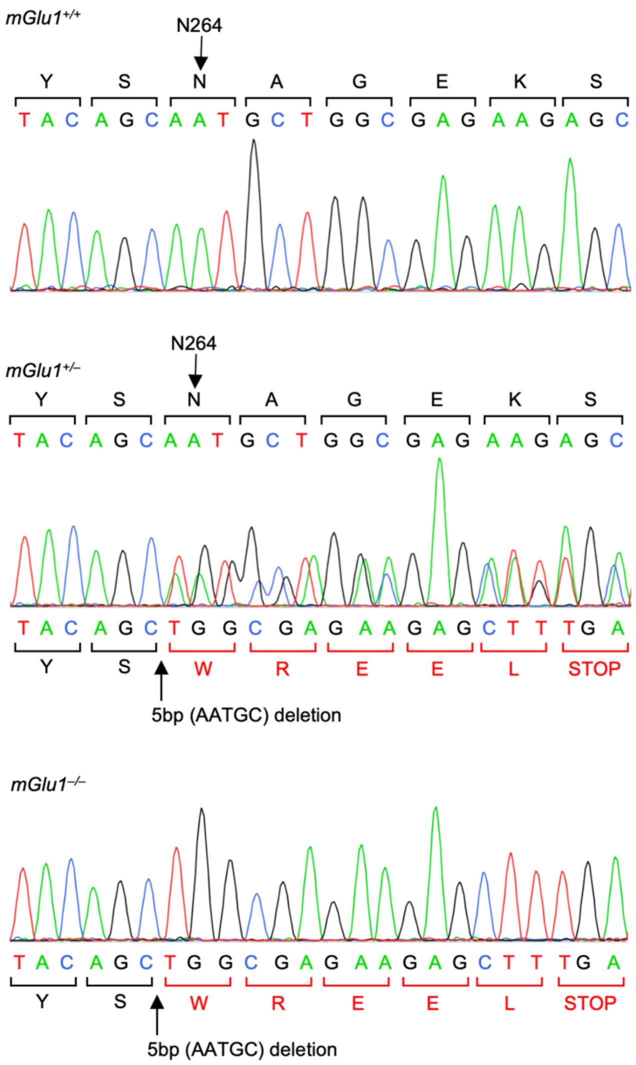


**Supplementary figure 12 | Sequence analyses of exon 3 of mGlu1 using *mGlu1*$^{+/+}$, *mGlu1*$^{+/–}$, or *mGlu1*$^{–/–}$ genome.**

### Supplementary Methods

#### Gross cerebellar anatomy

To observe gross structure in adult mouse cerebella, cresyl violet-staining was performed, as previously described[S1]. Bright-field images were captured using a CCD camera (DP70, Olympus) attached to a stereomicroscope (SMZ1000, Nikon).

#### Rota-rod test

To evaluate motor coordination of *mGlu1*$^{+/+}$, *mGlu1*$^{CBC/CBC}$ or *mGlu1*$^{-/-}$ mice, accelerating rota-rod test was performed at 4−40 rpm using a single lane rota-rod treadmill (MK-630B, Muromachi Kikai Co., Ltd.), and the time that each mouse stayed on the rod was measured.

### Synthesis and Characterization

#### General materials and methods for organic synthesis

All chemical reagents and solvents were purchased from commercial sources (FUJIFILM Wako pure chemical, TCI chemical, Sigma-Aldrich) and were used without further purification. Thin layer chromatography (TLC) was performed on silica gel 60 F254 precoated aluminum sheets (Merck). Chromatographic purification was performed using flash column chromatography on silica gel 60 N (neutral, 40–50 µm, Kanto Chemical). $^{1}$H-NMR or $^{13}$C-NMR spectra were recorded in deuterated solvents on an Advance III HD 500 MHz or Advance III HD 300 MHz (Bruker). Chemical shifts were referenced to residual solvent peaks or tetramethylsilane (δ = 0 ppm). Multiplicities are abbreviated as follows: s = singlet, d = doublet, t = triplet, m = multiplet, brs = broad singlet. High resolution mass spectra were measured on a compact (Bruker) equipped with electron spray ionization (ESI). Reversed-phase HPLC (RP-HPLC) was carried out on a Hitachi Chromaster system equipped with a UV detector, and an YMC-Pack ODS-A column.

### Synthesis of Compound FITM-Cy3

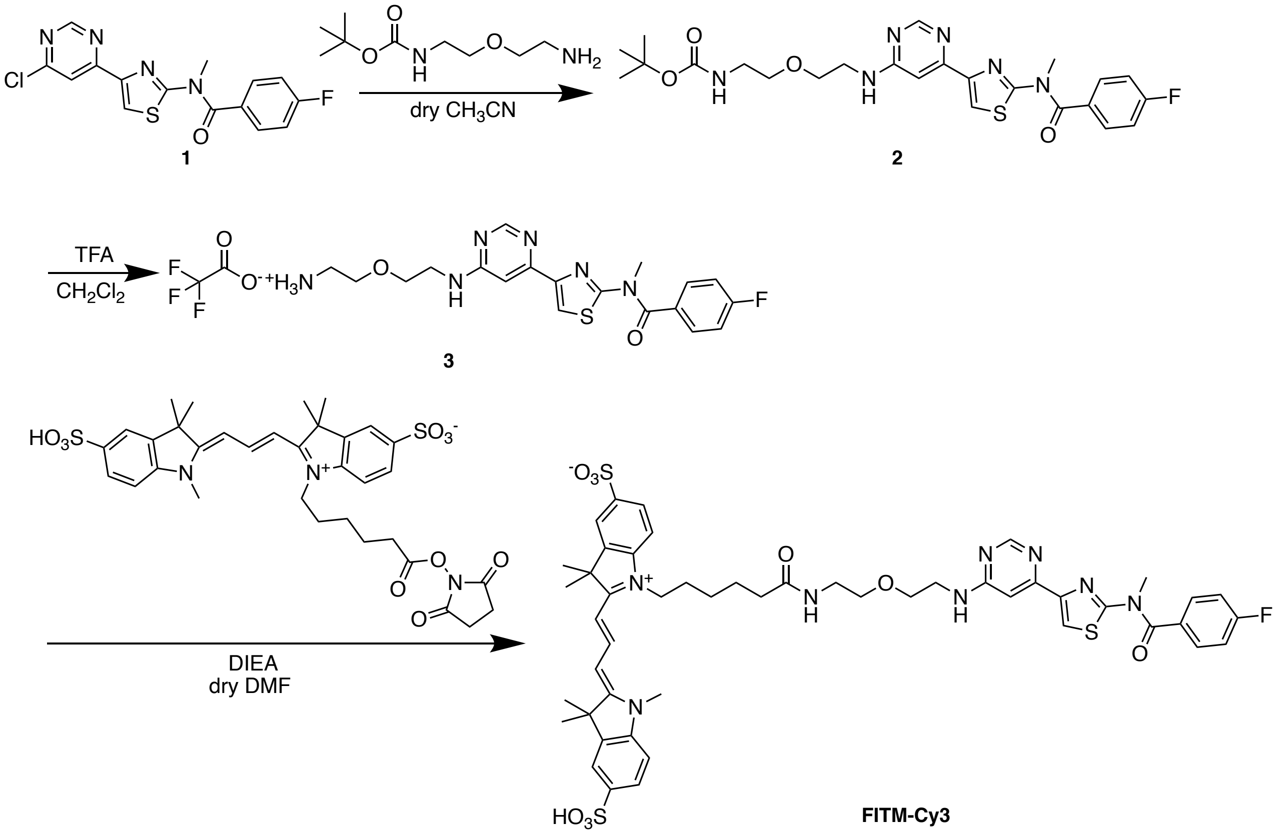


Synthesis of Compound **2**

*N*-(*tert*-Butoxycarbonyl)-2-(2-aminoethoxy) ethylamine (ref. S2) (50 mg, 0.25 mmol) was added to a solution of **1** (ref. S3) (30 mg, 86 μmol) in dry $CH_3CN$ (5 mL). The reaction mixture was stirred for 61 h in a $N_2$ atmosphere at 80 °C. The reaction mixture was evaporated to distil away $CH_3CN$, and $CHCl_3$ was added to the residue. The mixture was washed with saturated $NH_4Cl$ aqueous solution and brine. The organic layer was dried with $Na_2SO_4$ and concentrated *in vacuo*. The residue was purified by column chromatography (silica gel, EtOAc/n-hexane = 2/1) to afford compound **2** (33 mg, 0.064 mmol, 74% yield) as a colorless amorphous substance. $^1$H-NMR (500 MHz, $CDCl_3$) $\delta$ 8.59 (s, 1H), 7.93 (s, 1H), 7.61 (dd, $J$ = 5.3, 8.8 Hz, 2H), 7.20 (t, $J$ = 8.8 Hz, 2H), 7.13 (d, $J$ = 1.0 Hz, 1H), 5.54 (brs, 1H, NH), 4.97 (brs, 1H, NH), 3.74 (s, 3H), 3.68 (brt, $J$ = 4.2 Hz, 2H), 3.66-3.60 (m, 2H), 3.55 (brt, $J$ = 5.1 Hz, 2H), 3.36-3.30 (m, 2H), 1.44 (s, 9H); $^{13}$C-NMR (126 MHz, $CDCl_3$) $\delta$ 169.6, 165.3, 163.3, 160.5, 158.6, 156.2, 132.6, 131.0, 130.6, 130.3, 128.9, 116.1, 79.6, 70.4, 69.7, 68.3, 41.1, 40.5, 38.6, 28.5; HRMS (ESI$^+$): calcd for $[M + Na]^+$ ($C_{24}H_{29}FN_6NaO_4S$) 539.1847, found 539.1851.

Synthesis of **FITM-Cy3**

Trifluoroacetic acid (200 μL, 298 mg, 2.6 mmol) was added to a solution of **2** (28 mg, 54 μmol) in $CH_2Cl_2$ (1 mL). The reaction solution was stirred for 1 h at room temperature. After confirming the consumption of **2**, the reaction solution was concentrated *in vacuo* to obtain **3**. Sulfo-Cy3 NHS ester

(1 mg, 1.3 μmol) and *N*,*N*-diisopropylethylamine (5 μl, 3.7 mg, 28.7 μmol) were added to a solution of **3** (5.0 mg, 9.4 μmol) in dry DMF (500 μL). The reaction solution was stirred overnight in a $N_2$ atmosphere at room temperature. The residue was purified by RP-HPLC (ODS-A, 250 x 10 mm, mobile phase; $CH_3CN$: 10 mM $AcONH_4$ aq. = 5:95 (0 min), 65:35 (60 min), flow rate; 3.0 mL/min, detection; UV (220 nm)) to afford FITM-Cy3 (0.58 μmol determined using the molecular extinction coefficient of Sulfo-Cy3 NHS ester (162,000 $M^{-1}cm^{-1}$), 45% yield) as a magenta powder. $^1$H NMR (500 MHz, $CD_3OD$) $\delta$ 8.52 (t, *J* = 13.5 Hz, 1H), 8.38 (s, 1H), 7.95 (d, *J* = 1.6 Hz, 1H), 7.94 (d, *J* = 1.6 Hz, 1H), 7.91 (dd, *J* = 1.6, 8.3 Hz, 1H), 7.91 (dd, *J* = 1.6, 8.3 Hz, 1H), 7.89 (s, 1H), 7.68 (dd, *J* = 5.3, 8.8 Hz, 2H), 7.38 (d, *J* = 8.3 Hz, 1H), 7.34 (d, *J* = 8.3 Hz, 1H), 7.29 (t, *J* = 8.8 Hz, 2H), 7.28 (d, *J* = 0.9 Hz, 1H), 6.45 (d, *J* = 13.5 Hz, 1H), 6.40 (d, *J* = 13.4 Hz, 1H), 4.09 (brt, *J* = 7.5 Hz, 2H), 3.72 (s, 3H), 3.69 (s, 3H), 3.64 (brt, J = 4.5 Hz, 2H), 3.62-3.56 (m, 2H), 3.52 (t, *J* = 5.5 Hz, 2H), 3.34 (t, *J* = 5.5 Hz, 2H), 2.18 (brt, *J* = 6.4 Hz, 2H), 1.83-1.76 (m, 2H), 1.71-1.63 (m, 2H), 1.45-1.38 (m, 2H); HRMS (ESI$^-$): calcd for $[M - H]^-$ ($C_{49}H_{54}FN_8O_9S_3$) 1013.3165, found 1013.3163.

**Synthesis of Pd(sulfo-bpy)**

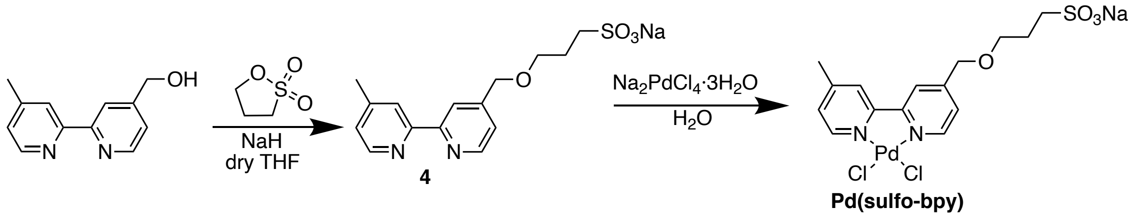


Synthesis of compound **4**

A solution of 4-hydroxymethyl-4'-methyl-2,2'-bipyridine (100 mg, 0.5 mmol) and sodium hydride (24 mg, 1.0 mmol) in dry THF (5 mL) was stirred on ice for 30 min under $N_2$ atmosphere. 1,3-propane sultone 73 mg (0.6 mmol) was added to the solution and stirred for 14 h at room temperature. After removal of the solvent by evaporation, the crude was purified by RP-HPLC (ODS-A, 250 x 10 mm, mobile phase; $CH_3CN$: 10 mM $AcONH_4$ aq. = 5:95 (0 min), 5:95 (10 min), 40:60 (60 min), flow rate; 3.0 mL/min, detection; UV (220 nm)) and neutrized with 1M NaOH, giving compound **4** (60 mg, 0.17 mmol, 34%) as a colorless oil. $^1$H-NMR (500 MHz, $CD_3OD$) $\delta$ 8.61 (d, *J* = 5.0 Hz, 1H), 8.51 (d, *J* = 5.0 Hz, 1H), 8.23 (s, 1H), 8.15 (s, 1H), 7.46 (d, *J* = 5.0 Hz, 1H), 7.34(d, *J* = 5.0 Hz, 1H), 4.66 (s, 2H), 3.71 (t, *J* = 6.3 Hz, 2H), 3.00-2.96 (m, 2H), 2.48 (s, 3H), 2.18-2.13 (m, 2H). $^{13}$C-NMR (126 MHz, $D_2O$) $\delta$ 155.8, 150.9, 150.4, 150.0, 148.7, 144.8, 126.6, 123.9, 120.6, 120.5, 70.3, 69.2, 47.8, 24.4, 21.0.

Synthesis of **Pd(sulfo-bpy)**

A solution of compound **4** (55 mg, 160 μmol) in $H_2O$ (3.0 mL) was added to a solution of sodium tetracholoropalladate trihydrate (II) (56 mg, 160 μmol) in $H_2O$ (2.0 mL). The solution was stirred at room temperature for 12 h. After concentration of the solution to 0.5 ml by evaporation, crystals were obtained from vapor diffusion of acetone into a $H_2O$ at room temperature, giving Pd(sulfo-bpy) (49 mg, 94 μmol, 59%) as an orange solid. $^1$H-NMR (500 MHz, $CD_3OD$) $\delta$ 9.10 (s, 1H), 9.00 (s, 1H), 8.39 (s, 1H), 8.36 (s, 1H), 7.63 (d, $J$ = 6.0 Hz, 1H), 7.52 (d, $J$ = 6.0 Hz, 1H), 4.76 (s, 2H), 3.78 (t, $J$ = 6.0 Hz, 2H), 2.99 (t, $J$ = 7.5 Hz, 2H), 2.20-2.14 (m, 2H). $^{13}$C-NMR (126 MHz, $D_2O$): $\delta$ 155.1, 154.9, 154.4, 154.2, 149.2, 148.5, 128.1, 125.2, 124.6, 121.6, 69.8, 69.6, 48.0, 24.5, 21.1.

### Synthesis of Pd(EG-bpy)

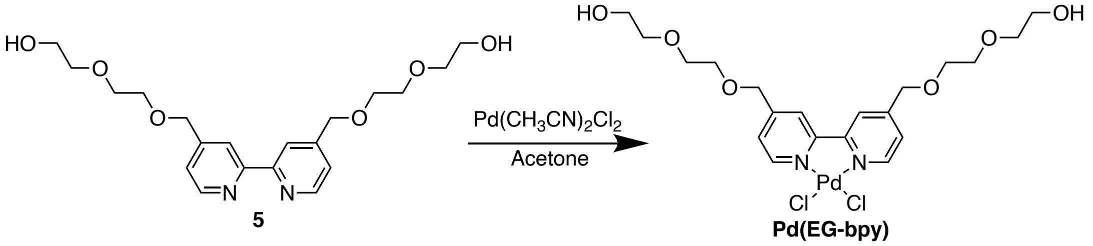


A solution of compound **5** (ref. S4) (10 mg, 26 μmol) in acetone (0.4 mL) was added to a solution of Bis(acetonitrile) dichloro palladiium (II) (6 mg, 23 μmol) in acetone. The solutioin was stood still at room temperature for 3 days, and filtered to obtain Pd(EG-bpy) (5.7 mg, 26 μmol, 59%) as an orange needle crystal. $^1$H-NMR (500 MHz, $CD_3OD$) $\delta$ 8.64 (d, $J$ = 5.0 Hz, 2H), 8.40 (s, 2H), 7.32 (m, 2H), 4.68 (s, 2H), 3.74 (t, J = 6.5 Hz, 4H), 2.98 (m, 4H), 2.17 (m, 4H); $^{13}$C-NMR (126 MHz, $CDCl_3$) $\delta$ 155.8, 153.8, 149.9, 124.1, 121.6, 72.7, 70.7, 70.5, 70.2, 61.6.

**Synthesis of Pd(di-sulfo-bpy)**

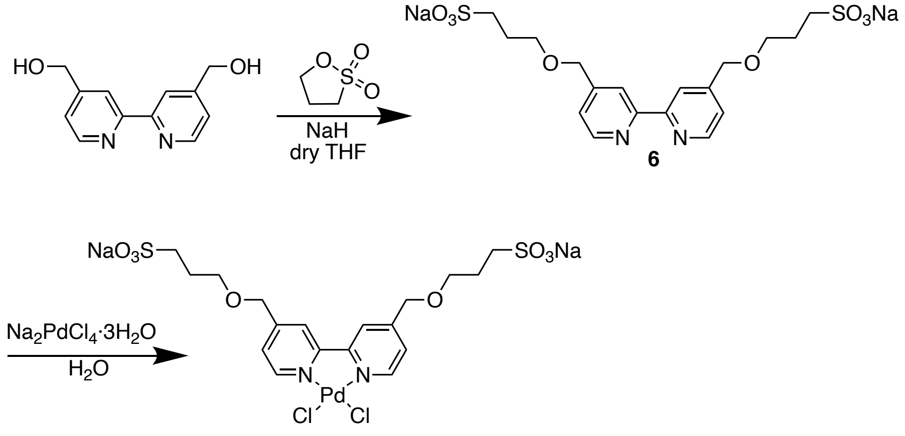

**Pd(di-sulfo-bpy)**

Synthesis of coumpound **6**

A solution of 4,4'-hydroxymethyl-2,2'-bipyridine (200 mg, 0.92 mmol) and sodium hydride (48 mg, 2.0 mmol) in dry THF (5 mL) was stirred on ice for 15 min under $N_2$ atmosphere. 1,3-propane sultone 244 mg (2.0 mmol) was added to the solution and stirred for 3 days at room temperature. The precipitation was collected and purified by RP-HPLC (ODS-A, 250 x 10 mm, mobile phase; $CH_3CN$: 10 mM $AcONH_4$ aq. = 0:100 (0 min), 0:100 (10 min), 40:60 (60 min), flow rate; 3.0 mL/min, detection; UV (220 nm)), and neutrized with 1M NaOH, giving compound **6** (60 mg, 0.17 mmol, 34%) as a colorless oil. $^1$H-NMR (500 MHz, $CD_3OD$) $\delta$ 8.63 (d, $J$ = 5.0 Hz, 2H), 8.29 (s, 2H), 7.48 (d, $J$ = 5.0 Hz, 2H), 4.69 (s, 2H), 3.74 (t, $J$ = 6.5 Hz, 4H), 2.98-2.96 (m, 4H), 2.20-2.14 (m, 4H); $^{13}$C-NMR (126 MHz, $D_2O$) $\delta$ 154.9, 149.3, 149.0, 122.8, 120.4, 70.6, 69.1, 47.9, 24.4.

Synthesis of **Pd(di-sulfo-bpy)**

A solution of compound **6** (45 mg, 89 μmol) in MeOH (2.2 mL) was added to a solution of sodium tetracholoropalladate trihydrate (II) (23 mg, 89 μmol) in MeOH (12 mL). The solution was stirred at room temperature for 1 h. After removal of the solvent by evaporation, the solid was dissolved in MeOH, and reprecipitated with diethylether, giving Pd(di-sulfo-bpy) (18 mg, 26 μmol, 30%) as an orange solid. $^1$H-NMR (300 MHz, $CD_3OD$) $\delta$ 9.14 (d, $J$ = 6.0 Hz, 2H), 8.40 (s, 2H), 7.70 (d, $J$ = 6.6 Hz, 2H), 4.77 (s, 2H), 3.77 (t, $J$ = 6.6 Hz, 4H), 3.00-2.95 (m, 4H), 2.20-2.11 (m, 4H); $^{13}$C-NMR (126 MHz, $D_2O$): $\delta$ 155.6, 154.2, 149.5, 124.8, 121.9, 69.9, 69.6, 48.0, 24.5.
